# Soft drinks and monogenetic diabetes: a study on the Wolfram syndrome 1 (*Wfs1*) deficient mouse model

**DOI:** 10.1101/318683

**Authors:** Rando Porosk, Julia Pintšuk, Marite Punapart, Ursel Soomets, Anton Terasmaa, Kalle Kilk

## Abstract

In a modern society, the risk of developing type II diabetes and obesity may be linked to the increased consumption of carbohydrate-rich drinks. Several genes, including Wolfram Syndrome 1 (WFS1), have been reported to increase susceptibility for developing type II diabetes. In this study we aimed to investigate the effect of chronic consumption of carbohydrate-rich drinks on weight gain, overall consumption of liquids, glucose tolerance and liver metabolism in *Wfs1*-deficient mice. *Wfs1*-deficient and wild-type mice were divided into three groups that consumed regular Coca-Cola, 20% sucrose solution or water *ad libitum* as the only source of liquid. During the experiment, daily liquid consumption was determined. After 30 days, total weight gain of mice was calculated and glucose tolerance test was performed. The liver tissue was analysed by means of untargeted and targeted metabolomics using liquid chromatography-mass spectrometry. Weight gain was strongly affected by mouse genotype (p<0.001), their drink (p<0.001) and the interaction of both genotype and drink (p<0.001). Coca-Cola significantly increased liquid consumption in knock-out mice. There was an effect of the drink (p<0.001) and the interaction between the genotype and treatment (p=0.02) on blood glucose level while Coca-Cola and 20% sucrose solution exacerbated glucose intolerance in the knock-out mice. In untargeted metabolic profiling, the water consuming wild-type and heterozygous mice were found to be the most distinctive from the mice with all other genotype and drink combinations. Targeted analysis revealed interactions between the genotype and drink regarding to glycolysis and lipogenesis. In the wild-type animals, carbohydrate overload was alleviated by converting glucose to lipids. However, the same mechanism is not implemented in knock-out animals, as lipolysis and gluconeogenesis are upregulated by *Wfs1* deficiency. In conclusion, our study demonstrates a significant interaction between the genotype and the drink when comparing wild-type and *Wfs1* knock-out mice consuming soft drinks.

## Introduction

One of the leading sources of added sugars is soft drinks and their regular consumption has been linked to metabolic syndrome, obesity and diabetes [1,2]. Habitual consumption of sweetened beverages may subsequently affect insulin resistance, carbohydrate and lipid metabolism and hepatic steatosis [3]. In rats, continuation of high-fructose diets has been shown to induce fatty liver, increased hepatic lipid peroxidation and activation of inflammatory pathways in the liver [4].

Type I and type II diabetes are generally polygenic in nature and may be caused by multiple environmental and genetic factors. On the other hand, diabetes can be monogenic (5-10% of cases), which is characterized as a heterogeneous group of rare disorders due to genetic mutations/defects in single genes, causing dysfunction of beta cells and hyperglycemia. An ongoing list of genes/loci has been associated with monogenic diabetes and insulin resistance [5], while hyperglycemia is one of the common consequences of mutations in these genes. Every single gene associated with monogenic diabetes expresses a distinct phenotype and clinical features. Genome-wide association studies have found that one of the genes associated with the development of type II diabetes is the Wolfram syndrome 1 gene (*WFS1*) [6].

Biallelic mutations of the *WFS1* gene (encoding glycoprotein wolframin) cause a rare autosomal recessive neurodegenerative disease, Wolfram syndrome 1. Wolframin is a transmembrane protein located in the endoplasmic reticulum (ER) and lack of its normal function leads to impairment in ER stress response and to induction of apoptosis [7–10]. ER stress is closely related to pathogenesis of diabetes as it alone can initiate beta cell failure and death in diabetes [11]. Homozygous mutations in the *WFS1* gene lead to the Wolfram syndrome, which is characterized by early-onset diabetes mellitus, progressive optic nerve atrophy, diabetes insipidus, and deafness [12,13].

We have previously generated and characterized *Wfs1*-deficient mice [14–16]. They exhibit impaired glucose tolerance due to insufficient insulin secretion [17] and wolframin may play an important role in maintaining homeostasis in beta-cells [18,19]. Female *Wfs1*-deficient mice tend to have lower blood glucose level and impaired glucose tolerance with relatively high insulin levels [20]. In addition, despite elevated growth hormone and insulin-like growth factor 1 levels, *Wfs1*-deficient mice have reduced body weight compared to their wild-type littermates [21]. Thus, deletion in *Wfs1* induces growth retardation although growth hormone production is activated [21]. The expression of phosphoinositide-3-kinase is reduced in the temporal lobe of *Wfs1*-deficient mice as revealed with the use of the Affymetrix gene chip array [21].

In this study Coca-Cola was chosen because it is a commercially available sucrose-sweetened soft drink worldwide and therefore highly standardisable. As WFS1 has been associated with the development of both type I and type II diabetes, the aim of this study was to evaluate the effect of the genotype and the drink of chronic consumption (30 days) of soft drinks on diabetes progression using wild-type and *Wfs1*-deficient mice as the study subjects.

## Materials and methods

### Drugs and chemicals

Refined white sugar (sucrose, food grade) was purchased at a local grocery store, and a 20% (w/vol) drinking solution was made, using tap water, and prepared prior to administration. Regular Coca-Cola containing sucrose 110 mg/ml (Coca-Cola HBC, Krakow, Poland) was purchased at the local grocery store and was degassed before pouring into animal drinking bottles. All other chemicals were purchased from Sigma-Aldrich (Merck KGaA, Darmstadt, Germany).

### Wfs1-deficient mice

The generation of *Wfs1*-deficient mice has been described elsewhere[21]. All studies were performed on female F2 hybrids [(129S6/SvEvTac x C57BL/6) x (129S6/SvEvTac x C57BL/6)] who were 5-6 months old at the time of testing. The mice were housed in groups of 8-9 at 20±2 °C under a 12-h/12-h light/dark cycle (lights on for 07:00 hours), with free access to food pellets and water. The animal experiments described in this study were performed with a permission from the Estonian National Board of Animal Experiments (No. 86, August 28, 2007) and in accordance with the European Community Directive (86/609/EEC).

### Experimental groups

The animals were divided randomly, based on the genotype, into nine groups (n = 8) according to the source of the liquid utilized. Female wild-type (*Wfs1^+/+^*, WT) and *Wfs1*-deficient (heterozygous *Wfs1^+/-^*, HZ and homozygous *Wfs1^-/-^*, KO) mice were used throughout this study. The wild-type, HZ and KO control animals ingested water, 20% sucrose solution or regular Coca-Cola *ad libitum* as the only source of liquid.

### Glucose tolerance test

The mice were kept in their home cage with free access to food and water. Food was removed and sweetened liquids were replaced by water 12 hours prior to the glucose tolerance test and access to food pellets was prevented during the glucose tolerance test. Basal levels of blood glucose were determined from the tail vein, thereafter glucose (2 g/kg, i.p.) was administered. Blood glucose levels were measured 30, 60, 120 and 180 minutes following glucose injection. Glucose concentration in the blood was determined using a hand held glucose meter (Accu-Check Go, Roche, Mannheim, Germany).

### Sample preparation

Mice were sacrificed by cervical dislocation and the liver was dissected, perfused with ice-cold saline, snap frozen in liquid nitrogen and stored at -80°C until processing. The protocol for homogenization of the liver, described previously by Beckonert, was used in a slightly modified version [22]. For hydrophilic and lipophilic extraction of metabolites, the frozen samples were weighed and 4 ml/g of LC-MS grade methanol and 0.85 ml/g of water were added before homogenization. The samples were homogenized by a ultrasound homogenizer (Bandelin Sonopuls, Berlin, Germany), followed by adding 4 ml/g of chloroform and 2 ml/g of water. The samples were mixed and centrifuged for 15 min 1000 × g at 4°C, which allowed the mixture to settle into two layers (the upper hydrophilic and the lower lipophilic phase). These layers of the tissue homogenates were then separated. The proteins from the separated layers were precipitated with 75% methanol and centrifuged for 15 min at 21250 × g 4°C. All procedures were carried out on ice.

### Mass-spectrometry

The samples were randomized and the supernatants were analyzed on a Q-Trap 3200 mass spectrometer (AB Sciex, Framingham, MA, USA). The samples were analyzed for 5 min in the isocratic flow of 0.05 ml/min of methanol and 0.1% of formic acid for the lipophilic phase and in the binary flow of 0.025 ml/min of water and methanol, followed by 0.05 ml/min of methanol and 0.1% formic acid for the hydrophilic phase. The full spectra (mass-to-charge ratio 50 to 1500) were obtained in the positive and negative enhanced mass scan mode. The ionspray voltage, and the declustering and entrance potentials were 4500 V, 20 V and 10 V, respectively, and the corresponding negative voltages were applied for the negative scan mode.

Along with metabolic profiling, targeted metabolomics was carried out with selected hydroxyl acids, all proteogenic amino acids and acylcarnitines using a previously described method[16]. Acylcarnitines (free, acetyl-, propionyl- and butyrylcarnitine) were analyzed as the precursors of m/z 85 ion and all amino acids were analyzed by multiple reaction monitoring scan. Ionization was performed at 4500 V and 400 °C, the declustering potential was set at 40 V and collision energy was set at 38 V. For analysis of hydroxy acid and hexoses (phospho-, bisphospho- and monohexoses, lactate, succinate, citrate and oxaloacetate), 20 μl liver homogenate was mixed with 40 μl (500 μM [^2^H_4_]succinic acid in methanol). The samples were centrifuged for 15 min at 10 000 × g and 20 μl were analyzed by liquid chromatography-mass spectrometry.

### Quantitative determination of triglycerides

For determination of the overall level of triglycerides in the liver, the tissues were homogenized by an ultrasound homogenizator in 0.1 M phosphate buffer (1:10 w/v; pH 7.4), followed by the use of commercially available Triglycerides-LQ assay kit (Spinreact, Girona, Spain). The results were measured using Tecan Sunrise spectrophotometer (Tecan Group, Männedorf, Switzerland).

### Statistical analysis

The data were compared by one-way or multiple-way analysis of variance (ANOVA) followed by the Tukey post hoc test. Fitness to normality was assessed by histograms and normal probability plots. Variances between the groups were similar and a *p-*value of <0.05 was considered statistically significant. No values were determined as outliers and excluded. The spectral signals of the untargeted metabolic profiles were binned with a m/z resolution of 1 and normalized to mean intensity of the spectra. Statistical analysis was performed using STATISTICA version 9 (StatSoft Ltd, Bedford, UK), GraphPadPrism version 5 software (GraphPad Software Inc, San Diego, CA, USA) and R version 3.2.2 (The R Foundation for Statistical Computing).

## Results

### Gain of body weight

The knock-out mice had lower average body weight than their littermates (KO 17.53 g; HZ 22.58 g; WT 22.29 g) [one way ANOVA F(2,68)=58.35, p<0.001]. Thirty-day consumption of a 20% sucrose solution and Coca-Cola resulted in increased body weight in the WT mice (Fig. 1). The sucrose solution did not induce weight gain in the HZ mice but potentiated weight loss in the KO mice. In contrast, Coca-Cola induced weight gain in the HZ mice but had no effect on the body weight of the KO mice. There was a significant effect of treatment [two-way ANOVA F(2,62)=19.2, p<0.001] and genotype [F(2,62)=54.8, p<0.001] as well as of the interaction between the two [F(4,62)=7.7, p<0.001].

**Fig 1.**
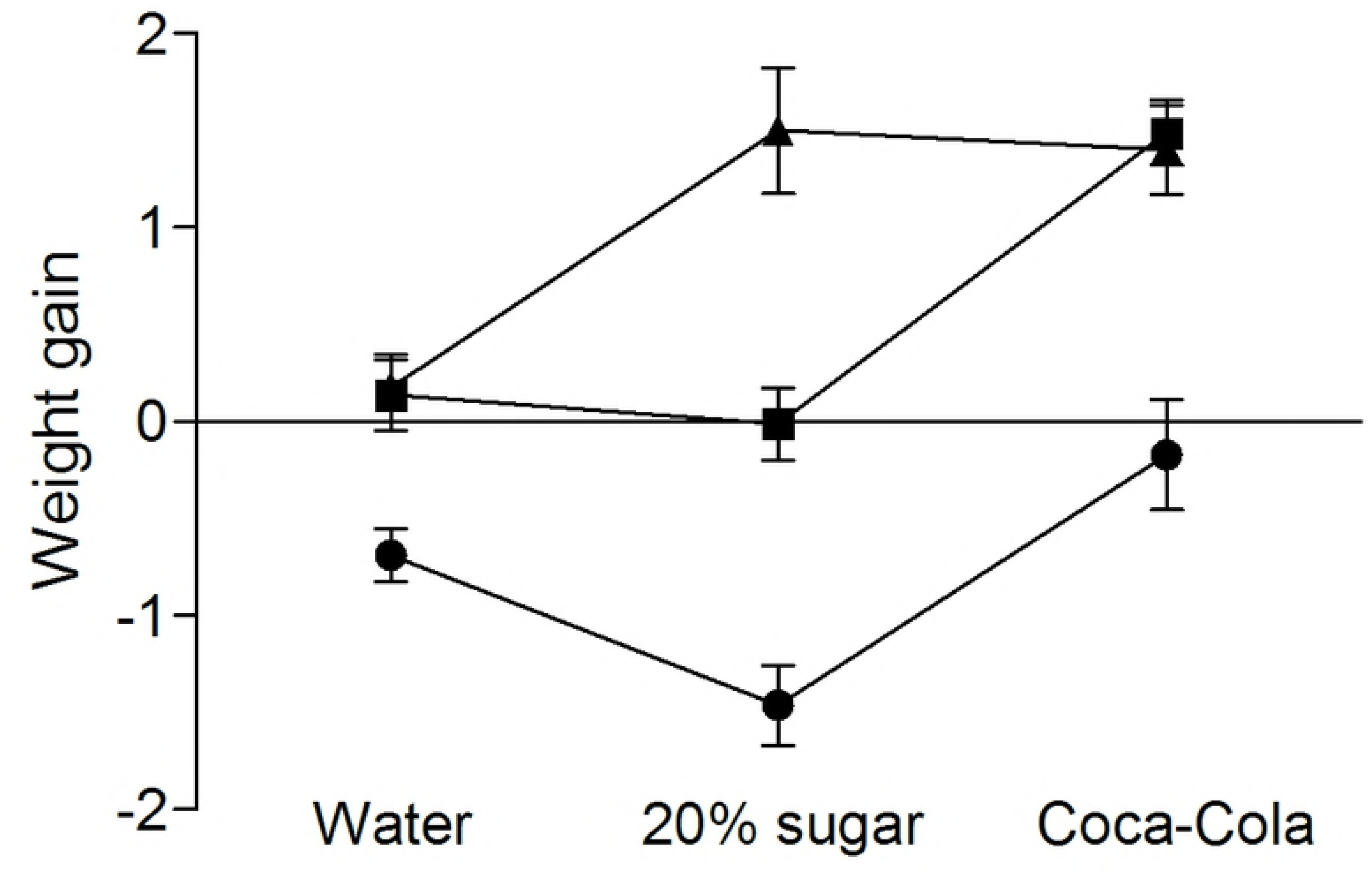
Body weight gain. Gain of body weight in *Wfs1* knock-out (KO, circle), heterozygous (HZ, square) and wild-type (WT, pyramid) mice after 30-day consumption of *ad libitum* water, 20% sucrose solution or regular Coca-Cola. Data are presented as mean ± SEM.

### Liquid consumption

Liquid consumption was recorded as total per group (one bottle per cage) and average daily liquid consumption per mice per day was calculated. The mice consumed similar amounts of water irrespective of the genotype (Fig. 2). 20% sucrose solution increased liquid consumption in all genotypes with the strongest effect in the KO mice. Coca-Cola consumption had a mild if any effect on liquid consumption in the WT and HZ mice, but greatly increased liquid consumption in the KO mice.

**Fig 2.**
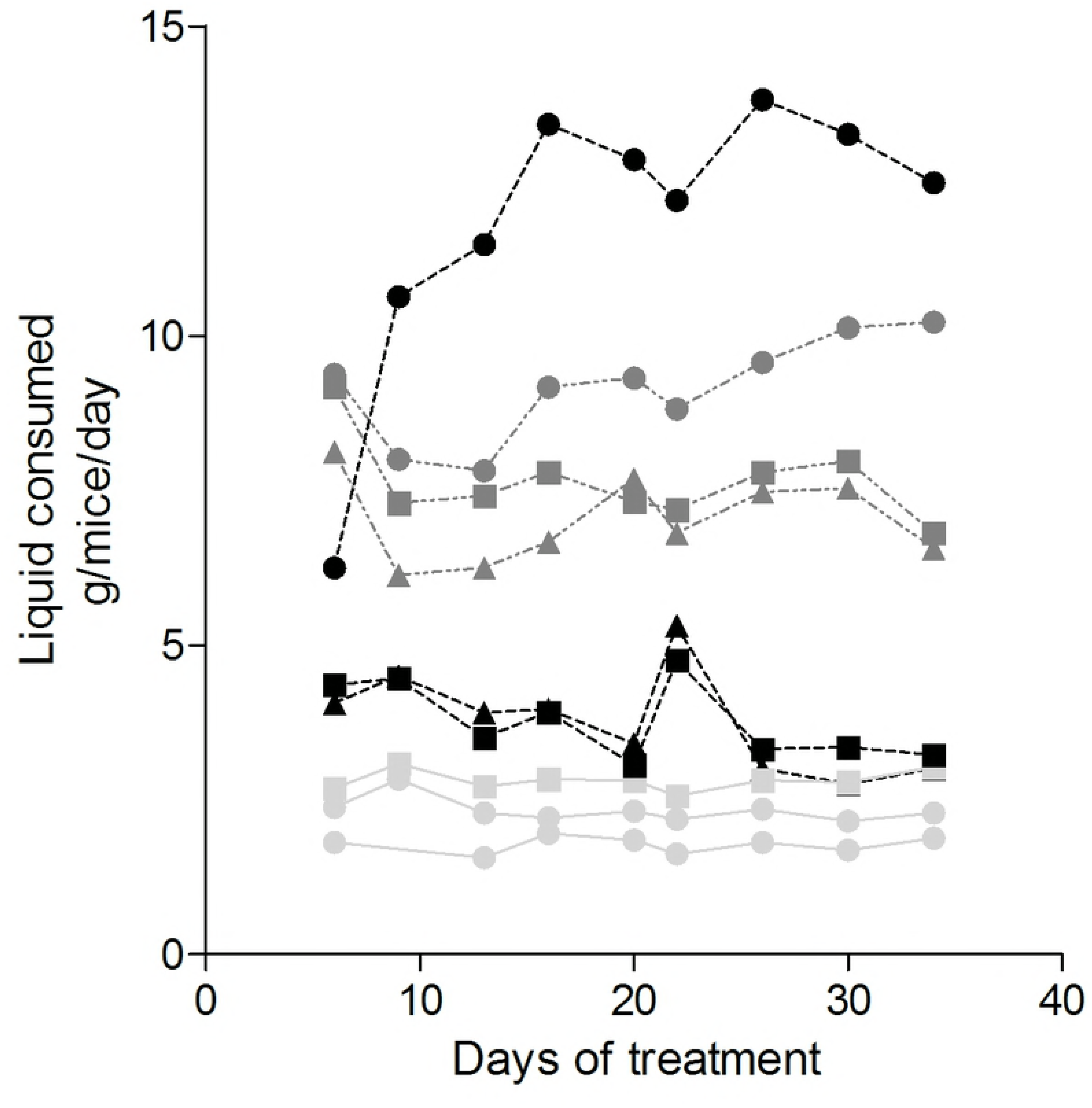
Liquid consumption. Daily liquid consumption in *Wfs1* knock-out (KO, circle), heterozygous (HZ, square) and wild-type (WT, pyramid) mice during 30-day consumption of *ad libitum* water (light grey), 20% sucrose solution (dark grey) or regular Coca-Cola (black).

### Blood glucose level

Blood glucose levels were measured at the end of the 30-day experiment. The mice were fasted for 12 hours before measurement of blood glucose level and the glucose tolerance test. Interestingly, basal blood glucose levels were not different across the genotypes (Fig. 3A). Treatment with 20% sucrose solution resulted in a decrease of basal blood glucose levels in all genotypes when compared to the water group while Coca-Cola treatment resulted in a similar decrease only in the KO mice. Thus there was a statistically significant effect of treatment [two-way ANOVA F(2,62)=12.3, p<0.001] and of the interaction between the genotype and treatment [F(4,62)=3.1, p<0.02].

**Fig 3.**
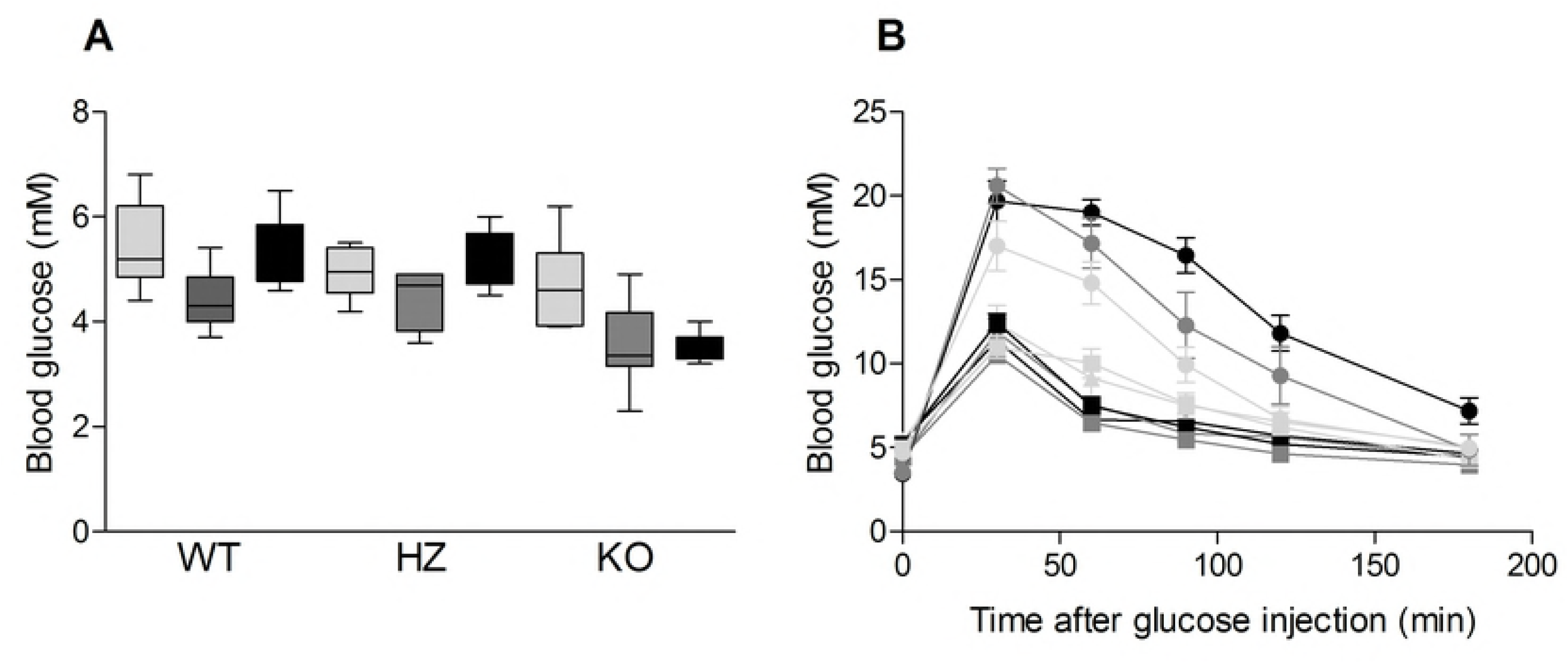
Glucose tolerance. The basal blood glucose level (A) and the intraperitoneal glucose tolerance test (B) of *Wfs1* knock-out (KO, circle), heterozygous (HZ, square) and wild-type (WT, pyramid) mice after 30-day consumption of *ad libitum* water (light grey), 20% sucrose solution (dark grey) or regular Coca-Cola (black), IPGTT 2 g/kg fasted. Data are presented as box and whisker plot showing median, 25^th^ and 75^th^ percentile, and min-max values.

### Glucose tolerance test

Administration of glucose (2 g/kg i.p.) induced a rise in blood glucose levels with a peak at 30 minutes in all genotypes (Fig. 3B). Blood glucose levels at 30 minutes after glucose administration were the highest in the KO mice. Coca-Cola or 20% sucrose solution had no significant effect on glucose tolerance in the WT mice. Coca-Cola treatment resulted in increased maximal blood glucose levels in the HZ mice when compared to the HZ water group. Surprisingly, glucose clearance was faster in the Coca-Cola HZ group when compared to the water group. The 20% sucrose solution did not produce a rise in maximal blood glucose levels but, similarly to Coca-Cola, improved the glucose clearance rate in the HZ mice. Coca-Cola and 20% sucrose solution exacerbated glucose intolerance in the KO mice.

### Metabolomics

Principal component analysis was performed to visualize variance in the liver metabolic profiles of the studied mice. The water consuming WT and HZ mice were partly separated from all other genotype and drink combinations which formed a single group (Fig. 4).

**Fig 4.**
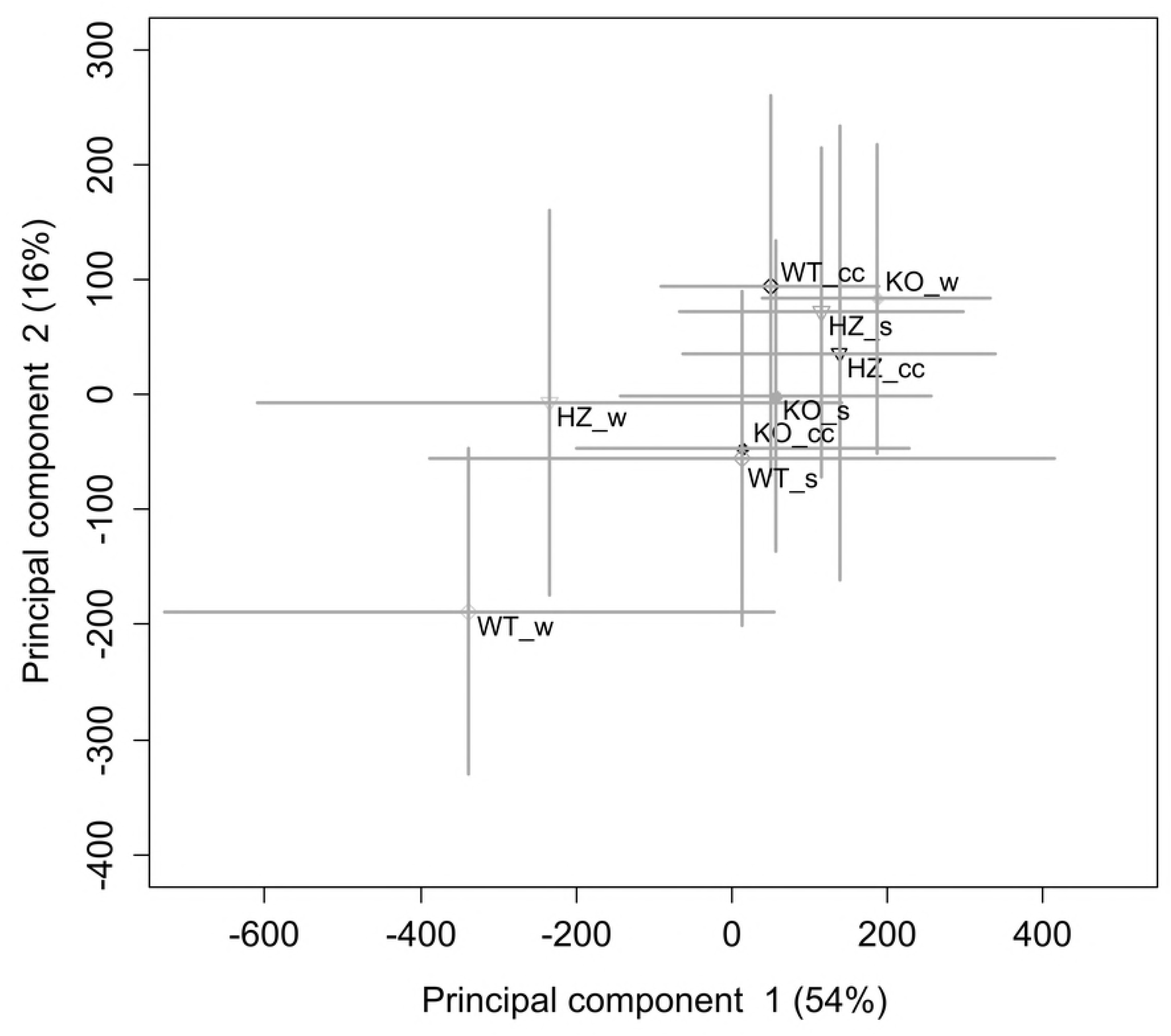
Principal component analysis of the liver metabolome. *Wfs1*-deficient knock-out (KO), heterozygous (HZ) and wild-type (WT) mice consuming water (w), 20% sucrose solution (s) or Coca-Cola (cc). The percentage of total variance that the component describes is presented in the parentheses following the axis titles. Each dot represents an array of metabolic profiles from the liver. Data are presented as mean ± SD.

Specific metabolites, including amino acids, acylcarnitines, total triglycerides and some glycolysis and citrate cycle intermediates, were quantified from the liver homogenate. The statistically significantly changed metabolites are presented in Table 1.

**Table 1.** Statistical differences in the liver tissue of *Wfs1*-deficient and wild-type mice due to genotype, diet or their interaction.

| Metabolite | Genotype |  |  | Diet |  |  | Interaction |  |  |
| --- | --- | --- | --- | --- | --- | --- | --- | --- | --- |
|  | df | F value | P value | df | F value | P value | df | F value | P value |
| Bisphosphohexoses | 2,60 | 0.35 | 0.70474 | 2,60 | 4.36 | **0.01708** | 4,60 | 0.97 | 0.43051 |
| Hexoses | 2,60 | 6.84 | **0.00211** | 2,60 | 0.80 | 0.45202 | 4,60 | 1.55 | 0.20025 |
| Lactate | 2,60 | 7.41 | **0.00134** | 2,60 | 1.19 | 0.31144 | 4,60 | 4.40 | **0.00348** |
| Oxaloacetate | 2,60 | 12.33 | **0.00003** | 2,60 | 0.33 | 0.72066 | 4,60 | 3.69 | **0.00944** |
| Phosphohexoses | 2,60 | 3.73 | **0.02985** | 2,60 | 4.54 | **0.01462** | 4,60 | 3.11 | **0.02153** |
| Ala | 2,60 | 0.14 | 0.86967 | 2,60 | 7.25 | **0.00152** | 4,60 | 2.64 | **0.04239** |
| Gln | 2,60 | 4.29 | **0.01820** | 2,60 | 6.63 | **0.00250** | 4,60 | 2.79 | **0.03414** |
| Glu | 2,60 | 9.52 | **0.00026** | 2,60 | 8.27 | **0.00067** | 4,60 | 1.00 | 0.41721 |
| His | 2,60 | 3.93 | **0.02486** | 2,60 | 2.75 | 0.07178 | 4,60 | 3.89 | **0.00711** |
| Lys | 2,60 | 1.34 | 0.26890 | 2,60 | 4.57 | **0.01425** | 4,60 | 3.45 | **0.01325** |
| Pro | 2,60 | 6.48 | **0.00283** | 2,60 | 5.52 | **0.00629** | 4,60 | 2.29 | 0.06964 |
| Ser | 2,60 | 0.55 | 0.58188 | 2,60 | 2.48 | 0.09259 | 4,60 | 4.52 | **0.00293** |
| Carnitine | 2,60 | 12.48 | **0.00003** | 2,60 | 4.54 | **0.01463** | 4,60 | 0.17 | 0.95085 |
| Triglycerides | 2,60 | 5.69 | **0.00546** | 2,60 | 1.06 | 0.35434 | 4,60 | 4.92 | **0.00170** |

## Discussion

Metabolism of carbohydrates depends both on their intake and genetic flexibility to regulate involved enzymes and other proteins. *WFS1* deficiency, either partial, as in the HZ genotypes, or total as in the KO genotypes, is a unique model where both type I and II diabetes can develop via endoplasmic reticulum stress. The present study focused on the effect of long-term consumption of Coca-Cola or carbohydrate-rich water for wild-type and *Wfs1*-deficient mice, with a specific attention on their weight gain, overall liquid consumption, blood glucose level, glucose tolerance and liver metabolome.

Consumption of hypertonic drinks increases the osmolality of blood. Via osmoreceptor activation, two mechanisms are employed to counter the osmolality increase: firstly, thirst for increased water intake, and secondly, antidiuretic hormone secretion for water loss reduction. Increased thirst is apparent in all genotypes when drinking a sucrose solution. We have previously shown that the concentration of hexoses in the urine and kidneys of KO mice is increased compared to their WT littermates [16]. Therefore, in KO animals hyperosmolar urine excretes the excessive amount of water, meaning that even higher water intake is required to compensate for the inefficiency of the kidneys. Thus, the highest sucrose solution consumption by KO animals is an expected result. In the current experiment the sucrose solution had higher carbohydrate concentration (731 mOsm/L) than Coca-Cola (493 mOsm/L) and its effect on thirst was therefore expected to be more pronounced. Previous studies have shown that mean liquid intake is significantly higher also in WT rats who drank regular Coca-Cola compared to their littermates who drank tap water [23,24]. We noted that the amount of Coca-Cola consumed by the WT and HZ mice was only slightly larger than the amount of water consumed by their littermates, however, a very drastic difference appeared for the KO animals. The effect of the genotype is particularly obvious when Coca-Cola consumption is compared to sucrose solution consumption: the KO mice drank more Coca-Cola, while the WT and HZ mice drank, as expected, less Coca-Cola and more sucrose solution. Caffeine, present in Coca-Cola but not in sucrose water, is usually considered to have a diuretic rather than an antidiuretic effect, and therefore its consumption should increase water loss and enhance thirst. Caffeine is a structural analogue of uric acid. We have previously shown that uric acid metabolism is severely affected by *Wfs1*-deficiency [16]. Additionally, in humans, sucrose containing soft drinks have been shown to interfere with uric acid production [25,26]. Uric acid in turn can enhance sucrose and fructose caused dysregulation in saccharide and lipid metabolism [26]. The anomalously high thirst in *Wfs1* KO Coca-Cola drinking animals may therefore be due to the interaction of the genotype with caffeine, uric acid and saccharide metabolism.

Increased consumption of sweetened beverages increased the body weight of the WT mice. Similar results have been reported by other authors: drinking regular Coca-Cola leads to increased body mass, usually in spite of a net decrease in solid food consumption [24]. The *Wfs1* HZ mice consumed of the same quantity sweetened beverages as the WT mice, but their body weight increased only when they were drinking Coca-Cola. Compared to the other groups, the KO animals consumed sweetened drinks the most but had the least increase in body weight. In contrary, weight loss was seen in the sucrose solution drinking KO animals. It has been previously shown that the KO mice consume less food when on standard diet compared to the WT littermates [27]. Additionally, it is known that the physical growth of the *Wfs1* deficient mice is retarded [21] and in a recent metabolomics study we hypothesized that defective collagen synthesis for connective tissue may be a key process [16]. Reduced solid food intake and reduced capability for tissue expansion both can counter the increased calories from drink. Coca-Cola, however, still causes weight gain. A recent study on humans has suggested a synergistic effect of caffeine and high carbohydrate intake [28], but there is more evidence that caffeine consumption is protective against obesity and diabetes [29]. Interestingly, several reports show caffeine efficacy in countering ER stress [30,31]. The latter being the main pathophysiology of Wolfram syndrome suggest a possible protective role of caffeine against *Wfs1* deficiency, which in turn allows body mass increase by excessive sugar intake. In summary, Coca-Cola consumption was associated with body weight increase in all genotypes, but *Wfs1* deficiency outweighed the effect of weight gain of the sucrose solution.

The blood glucose level decreased after the long-term consumption of 20% sucrose solution in all genotypes. This is not expected as high sucrose diet or soft-drink are usually reported to increase or not affect fasting blood glucose [32–34]. However, chronic consumption of high sucrose diet has been found to decrease fasting blood glucose in mice before, possibly due to altered insulin release kinetics [35]. The exact mechanism of this controversial outcome in our study is unknown, but the reduction in fasting blood glucose is in good correlation with the total sucrose consumed. The sucrose solution is more concentrated and consumed at higher quantities than Coca-Cola. Only the *Wfs1*-deficient KO mice drank more Coca-Cola and also had a more significant change in their blood sugar level.

The glucose tolerance test did not reveal any effect on the HZ or WT mice, which is consistent with the results of previous studies [23,36]. As the KO animals have impaired glucose tolerance then high carbohydrate drinks slightly worsen its degree. The glucose intolerance of the KO mice could partly explain the higher thirst in these animals but not the differences in the consumption of sucrose solution and Coca-Cola.

The metabolic profile of the liver in mass spectrometry demonstrates that the highest variance (i.e. the first principal component) partly differentiates the WT and HZ water drinkers from the others. Based on this very robust analysis, all mice kept on sucrose containing drinks and the water drinking KO mice had a relatively similar liver metabolite profile. Hence, although glucose intolerance can be observed only in KO genotype, the liver metabolism of the WT and HZ animals kept on carbohydrate-rich liquid has actually common traits with the effect of gene knock-out.

Since the metabolic profile provides little information about ongoing precise biochemical processes, a targeted analysis of specific compounds was also carried out. Liver metabolite analysis revealed a statistically significant effect of the genotype, drink, or their interaction in many glucose-related metabolites. While fasting blood sugar was decreased in all sucrose and KO Coca-Cola drinkers (Fig. 3A), the liver of the latter showed elevated levels of hexoses, lactate, alanine and serine (Fig. 5). Lactate is formed from the glycolysis end-product pyruvate if there is currently not enough capacity to direct pyruvate into the citric acid cycle and oxidative degradation. In the liver, lactate can, alternatively to catabolism, be used in gluconeogenesis for glucose re-synthesis. Its accumulation indicates that lactate production overwhelms all its elimination pathways. High level of glucose will supress gluconeogenesis. Glycolysis occurs, but in the case of the overwhelming citric acid cycle’s capacity to accept pyruvate, the formed pyruvate is converted to lactate via redox reactions and to alanine via transamination. Moreover, increased levels of 3-phosphoglycerate, an intermediate of glycolysis, may cause a rise in serine levels. In an earlier study of the *Wfs1* genotype, we found evidence that the *Wfs1* KO mice had increased gluconeogenesis [16]. In agreement with that study, oxaloacetate levels in the *Wfs1*-deficient mice were decreased in the current study. Also, there is a noteworthy interaction between the genotype and the drink: in WT animals carbohydrate drinks tend to decrease oxaloacetate levels, possibly via increased usage of oxaloacetate for citrate formation. In HZ and KO animals, on the contrary, a high carbohydrate drink leads to an increase in oxaloacetate. If the genotype-associated decrease in oxaloacetate were due to its use for gluconeogenesis, then a glucose rich drink should indeed supress the decrease. The absence of the increase in oxaloacetate in WT can be understood when one looks at triglyceride levels. A significant increase in triglyceride level suggests that excessive saccharides are converted to fats via glycolysis and lipogenesis. In WT rodents and other mammals, an increased concentration of triglycerides and a mild to severe fatty liver after consumption of high-carbohydrate drinks has been shown [1,23,24,37]. This process is evidently not effective in *Wfs1*-deficient animals.

**Fig 5.**
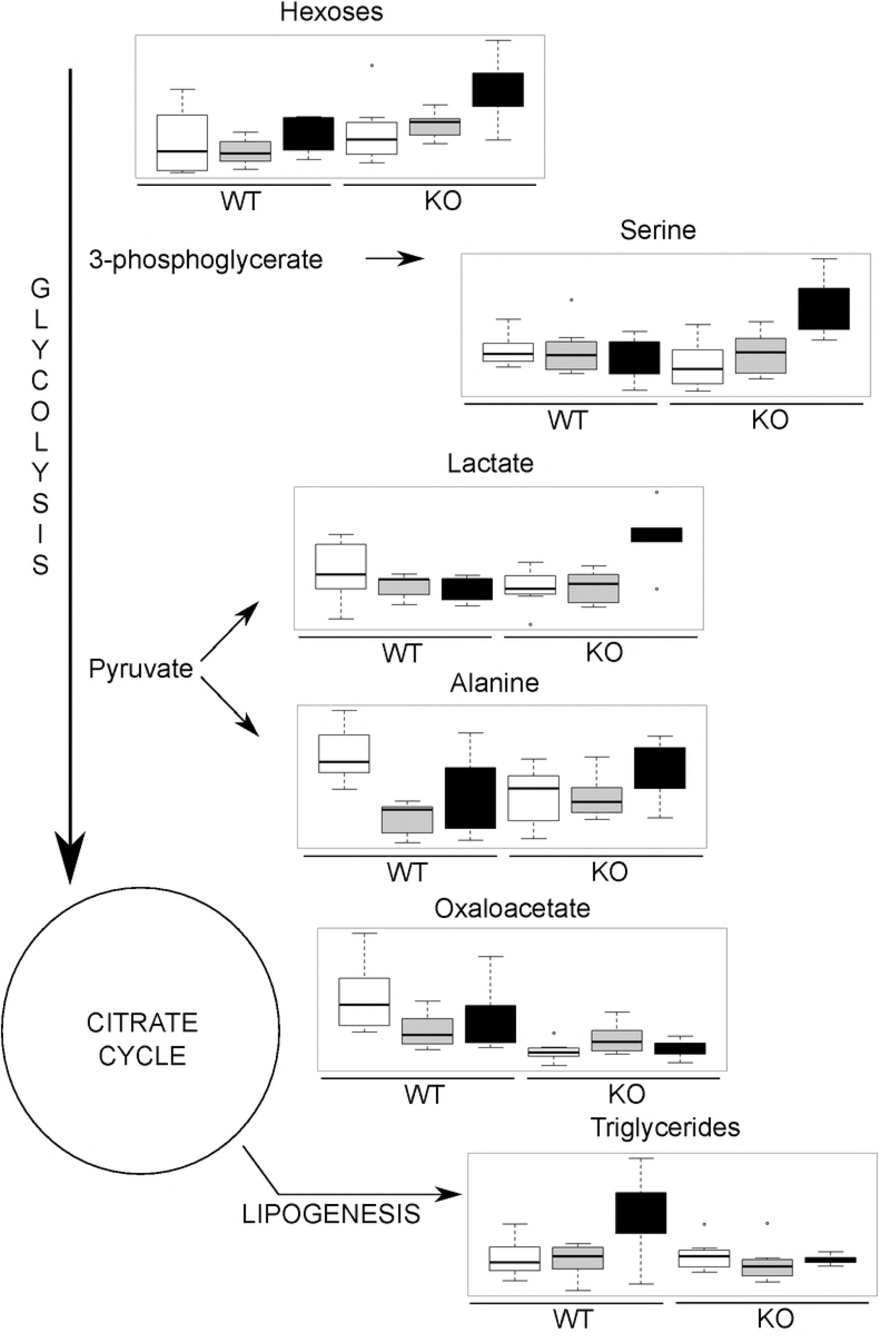
A simplified overview of the metabolic differences in *Wfs1* deficiency and diet. Shown are the glycolysis-related intermediates in the liver of wild-type (WT) and *Wfs1*-deficient (KO) mice consuming water (white), 20% sucrose solution (grey) or Coca-Cola (black). Data are presented as box and whisker plot showing median, 25^th^ and 75^th^ percentile, and min-max values.

When interpreting the results of liver metabolism one has to pay attention to the fact that when in the previous paragraph a carbohydrate “overload” mechanism was proposed, the described changes in metabolites were only apparent in the mice drinking Coca-Cola but not in those drinking the sucrose solution. The primary reason why this phenomenon appears only in this particular genotype and drink combination is liquid consumption. Although Coca-Cola has lower sucrose content than the 20% sucrose solution, the total carbohydrate load is the highest because of more intense thirst. Second, it cannot be excluded that caffeine or any other Coca-Cola ingredient has a crucial role. Caffeine is known to modulate gluconeogenesis in the liver [38,39], and although this effect cannot explain our results, some other effect may be involved. *Wfs1* deficiency is related to endoplasmic reticulum stress, which may alter enzyme sensitivities to xenobiotics.

Insulin resistance in peripheral tissues can enhance protein catabolism and amino acid usage for energetic needs. Compared to WT water drinkers, alanine level was decreased in most groups but KO Coca-Cola drinkers. In cases where a saccharide challenge is present, a decrease in amino acid levels can occur [40]. Increased levels of alanine and glutamine in the liver could indicate a particularly excessive catabolism of proteins (amino acids) in *Wfs1*-deficient Coca-Cola drinkers. During degradation of amino acids, the resulting nitrogen has to be transaminated to pyruvate to form alanine, and transported to the liver, activating the urea cycle and gluconeogenesis. We have previously described a decrease in most of the urea cycle intermediates in KO mice [16]. High-carbohydrate drinks did not seem to have a significant effect on the activity of the urea cycle. It may be though that the carbon overload from carbohydrates inhibits amino acid entry into the citrate cycle or even enhances reverse processes like α-ketoglutarate transamination to glutamate. Increased level of certain amino acids, glutamine in particular, is then not due to altered nitrogen or protein metabolism but yet another mode to store short chain carbon units.

Free carnitine level was decreased in both *Wfs1*-deficient genotypes and Coca-Cola decreased it further. Propionylcarnitine was increased in the KO mice who consumed Coca-Cola compared to the mice who drank water. This indicates higher usage of short-chain acylcarnitines for energy demand, which are shown to be altered in the serum in pre-diabetic conditions [41].

Taken together, the results of metabolomics indicate an interaction between the genotype and the drink in the metabolism of carbohydrates, which leads to increased levels of glycolysis intermediates and related metabolites. In WT animals, stress on the metabolism of carbohydrates is alleviated by converting them to lipids, however, as lipolysis and gluconeogenesis are upregulated by *Wfs1* deficiency, this mechanism does not work in KO animals. We have previously shown [16] that protein metabolism is genotype dependent. Based on this study, we suggest that the observed effect of the drink on amino acids originates from carbohydrate metabolism rather than from the direct influence of high carbohydrate drinks on protein metabolism.

Our study demonstrates that consumption of Coca-Cola does not increase the body weight of *Wfs1* KO mice but increases significantly their daily liquid consumption. Contrary to WT animals, KO animals do not convert excessive carbohydrates to triglycerides and their liver is overwhelmed by glycolysis intermediates and related metabolites. While soft drinks cause a fatty liver in WT animals, glucose intolerance is associated only with the genotype and not with the drink.

## Acknowledgements

This research was supported by Institutional Research Funding (grant no. IUT20-42, PUT1416), The Center of Excellence for Genomics and Translational Medicine from the Estonian Ministry of Education and Science and by the European Union through the European Regional Development Fund (project no. 2014-2020.4.01.15-0012).

